# To be browsed or not to be browsed: differences in nutritional characteristics of blackthorn *Prunus spinosa* subject to the long-term pressure of herbivores

**DOI:** 10.1101/2024.04.04.588043

**Authors:** Veronica Facciolati, Marcin Zarek, Ewa Błońska, Jarosław Lasota, Olga Orman, Michał Ciach

## Abstract

The impact of ungulates on temperate forest vegetation has been investigated for a long time. Numerous studies on food selection have identified the palatable plant species preferred by large European herbivores. However, intra-specific food selection and the question why particular plants of a given species are ignored during foraging have been neglected in the literature. In central Europe, Blackthorns *Prunus spinosa* growing in abandoned pastures are an important component of the red deer’s *Cervus elaphus* diet. In areas densely populated by deer, annual shoot browsing produces dwarf shrubby forms of blackthorns. However, some blackthorns are not browsed by ungulates and tend to adopt a tree-like form. The existence of distinct, browsing-dependent growth forms of blackthorns raises the question of inter-individual differences in the nutritional composition of plants. Based on factor analysis, we discovered differences in nutritional composition between browsed and unbrowsed blackthorns that might explain the individual plant-related drivers of red deer food preferences. The leaves of browsed blackthorns contained higher concentrations of C, N, P and Cu but lower levels of Ca and Mg than unbrowsed ones. Moreover, browsed blackthorns had a higher water content and higher concentrations of insoluble proteins, chlorophylls and carotenoids. We highlight the fact that the nutritional characteristics of an individual plant may explain the observed food selection pattern, leading to the unhindered growth of a fraction of the blackthorn population, in spite of severe pressure on the part of ungulate herbivores. The results of this study underline the important role of herbivores in the dynamics of plant communities, in which ungulates may mediate the persistence of certain individuals of a given species.

**Highlights:**

- Browsing by ungulates leads to the formation of dwarf shrubby forms in blackthorns
- Some blackthorns are unbrowsed and adopt a tree-like form
- Browsed blackthorns differ in chemical composition from unbrowsed ones
- The nutritional profile of a given plant may influence food selection by ungulates

## 1. Introduction

Mammalian herbivores tend to search for palatable and nutrient-rich food resources in order to satisfy their energetic and nutritional needs (Scogings et al. 2011; Jensen et al. 2015; Mahieu et al. 2021). Foraging selectivity among herbivores is mediated by multiple ecological and environmental drivers, including various defence strategies that plants employ (Perkovich and Ward 2022). The defensive reactions of plants can be classified in accordance with the desired result, i.e. resistance, tolerance and avoidance (Boege and Marquis 2005; Gómez et al. 2008; Lindroth and St. Clair 2013; Norghauer et al. 2014; Call and St. Clair, 2018). Firstly, resistance is the active means that plants use to withstand herbivore pressure by producing inducible defences. Secondly, in the case of tolerance, plants adapt in order to tolerate a certain amount of induced stress, for example, by induced resource sequestration (Orians et al. 2011; Scogings et al. 2014; Peschiutta et al. 2018). Thirdly, avoidance is a strategy aimed at minimizing damage, for example, by escaping browsing pressure by means of a boost of vertical growth, a phenomenon known as growth response (Call and St. Clair 2018). Not being mutually exclusive, the resistance and tolerance strategies can be used by plants simultaneously (Gordon and Prins 2008); plant defence strategies against browsing are therefore complex. As differences in environmental conditions may lead to a trade-off between plant growth and plant defence, herbivory may also contribute to the evolution of habitat specialization in plants (Grover and Holt 1998; Fine et al. 2006).

In a scenario where browsing is intense, individual plants can react by increasing the secretion of defensive compounds (Wigley et al. 2015), developing structurally induced defences (Salek et al. 2019), increasing foliar nitrogen (N) levels (Scogings et al. 2011) or changing resource allocation patterns (Orians et al. 2011; Scogings et al. 2014; Peschiutta et al. 2018). As plant defence systems may limit foraging preferences, ungulates have developed a number of strategies to counteract such systems, like avoidance behaviour, reducing the intake of certain food types (Iason and Villalba 2006) or physiological adaptation to tolerate toxic plant products with the aid of specific binding enzymes (Clauss et al. 2003; Iason and Villalba 2006; Mlambo et al. 2016; Perkovich and Ward 2022). Although plant morphology is suggested to have a greater influence on food preferences than the chemical composition of the browsed individuals (Hartley et al. 1997), there are conflicting results regarding the role of morphology as the principal driver (Rooke et al. 2004). Holeski et al. (2016) suggest that the foliage composition, comprising soluble carbohydrates, proteins and minerals, can affect ungulate dietary preferences. More specifically, concentrations of N, calcium (Ca), phosphorus (P), magnesium (Mg), copper (Cu) and sulphur (S) are known to play an important role in deer dietary preferences (Suttle 2010) and influence the selection of different plant species present in a given area (Ceacero et al. 2015).

Rosaceae, Fagaceae and Rubiaceae are among the preferred families of plants consumed by ungulates (Gebert and Verheyden-Tixier 2001; Storms et al. 2008; Freschi et al. 2021). The genus *Prunus* L. (Rosaceae) includes over 200 species of trees and shrubs (Arnold Arboretum, 1940), some of which are commercially important (Veličković et al. 2021), are used in the restoration of historical landscapes (Mijnsbrugge et al. 2016; Mijnsbrugge et al. 2022) and boost species diversity by providing appropriate foraging and habitat resources for wildlife (Popescu and Caudullo 2016). A general characteristic of plants of the genus *Prunus* is that they contain carbon (C)- and N-rich defensive compounds (Santos Pimenta et al. 2014; Peschiutta et al. 2018), and also Ca-based inducible defence mechanisms such as calcium oxalate crystals (COC) (Peschiutta et al. 2020).

Blackthorn *Prunus spinosa* is one of the most widespread and common wild shrub species in Europe (Mijnsbrugge et al. 2016), occurring mainly in central and western parts of the continent with range limits in southern Scandinavia, Asia Minor, Caucasus and North Africa (Seneta and Dolatowski 1997; Popescu and Caudullo, 2016). The species is commonly found along the margins of deciduous forests, as a component of thermophilous communities, and as one of the successional species along former meadows and arable fields (Szafer and Zarzycki 1977; Seneta and Dolatowski 1997; Popescu and Caudullo 2016). Blackthorn is insect-pollinated, and the seeds are commonly dispersed by birds and mammals that eat the fruits (Popescu and Caudullo 2016), although the species can also propagate by means of vegetative root suckers (Leinemann et al. 2014; Brown et al. 2022). Blackthorns have a higher level of morphological variability (Kobendza 1955; Browicz and Zieliński 1975; Hanelt 1997; Kosina 2023) and a larger inter-population heterogeneity than other woody species (Mijnsbrugge et al. 2013; Leinemann et al. 2014; Mijnsbrugge et al. 2016). Although there are not many studies on the growth and phenology of blackthorns (Mijnsbrugge et al. 2022), Leinemann et al. (2014) suggest that vegetative propagation and the long-distance zoo-dispersal of seeds are possible reasons for its considerable variability.

Blackthorn is listed among the woody species preferred by foraging red deer *Cervus elaphus* during summer and autumn (Dumont et al. 2005; Salek et al. 2019). It commonly grows on ecotones and abandoned farmland (Szafer and Zarzycki 1977), where it usually adopts the shrubby form (Seneta and Dolatowski 1997) or develops a dwarf form, apparently in response to long-term, high browsing pressure. However, despite ungulate foraging activity, some blackthorn individuals grow relatively high, reaching 4 m (Seneta and Dolatowski 1997) or even 6-8 m as tall tree-like forms (Kobendza 1955). This is the outcome of differential browsing pressure: heavily browsed trees assume a shrubby/dwarf form, whereas less or unbrowsed individuals may grow into a tree-like form (see Vourc’h et al. 2002 on western red cedar *Thuja plicata*; Kupferschmid et al. 2015 on silver fir *Abies alba*).

The aims of this study were to explore the chemical characteristics of individual blackthorns subjected to the long-term pressure of ungulates and to compare them with a group of plants that for reasons unknown were not browsed. We investigated the chemical profiles of the two groups of plants, expecting to discover differences between browsed and unbrowsed blackthorns with respect to the contents of elements responsible for plant growth or defence and nutrients known to be important drivers of food selection in ungulates.

## 2. Materials and methods

### 2.1. Study area

The study area was situated in the Western Bieszczady Mountains (SE Poland), which are part of the Carpathians, one of the largest mountain ranges in Europe. These mountains consist of sandstones and slates, and cambisols are dominant. The mean annual temperature is ca 8°C, July is the warmest month (mean air temperature 17°C) and January is the coldest one (–3°C). The mean annual precipitation is ca 850 mm, with the highest amounts falling in July (ca 120 mm) and the least in January (ca 40 mm). The growing season lasts about 225 days, and continuous snow cover persists for ca 80 days on average. Located in uniform soil conditions that developed on flysch formations (sandstone and shale), the study site was dominated by eutric cambisols with a silt loam grain size (the average content of sand was 20%, silt 73% and clay 7%) and a mull humus type, characterized by an efficient degree of decomposition (C/N ratio = ca 10).

For the purposes of this study, a complex of open habitats was selected, partly overgrown with tree and shrub thickets, representing the early successional stage of forest encroaching onto farmland, i.e. former arable fields and pastures (Fig. 1A). The study site was dominated by blackthorn, accounting for ca 80% of the entire coverage of trees and shrubs, along with hawthorns *Crataegus* spp. (10%), common pear *Pyrus communis* (5%), crab apple *Malus sylvestris* (3%) and roses *Rosa* spp. (2%). The majority of the blackthorn bushes found in the study area (ca 90%) betrayed signs of long-term browsing and had developed a shrubby form. Such individuals reached a maximum of 1 m in height, and showed traces of repeated shearing of the top and side shoots (Fig. 1B). In contrast, ca 10% of the blackthorn population in the study area were ca 3-5 m high, on which traces of deer browsing were negligible, usually limited to the tops of a few shoots (Fig. 1D).

**Fig. 1.**
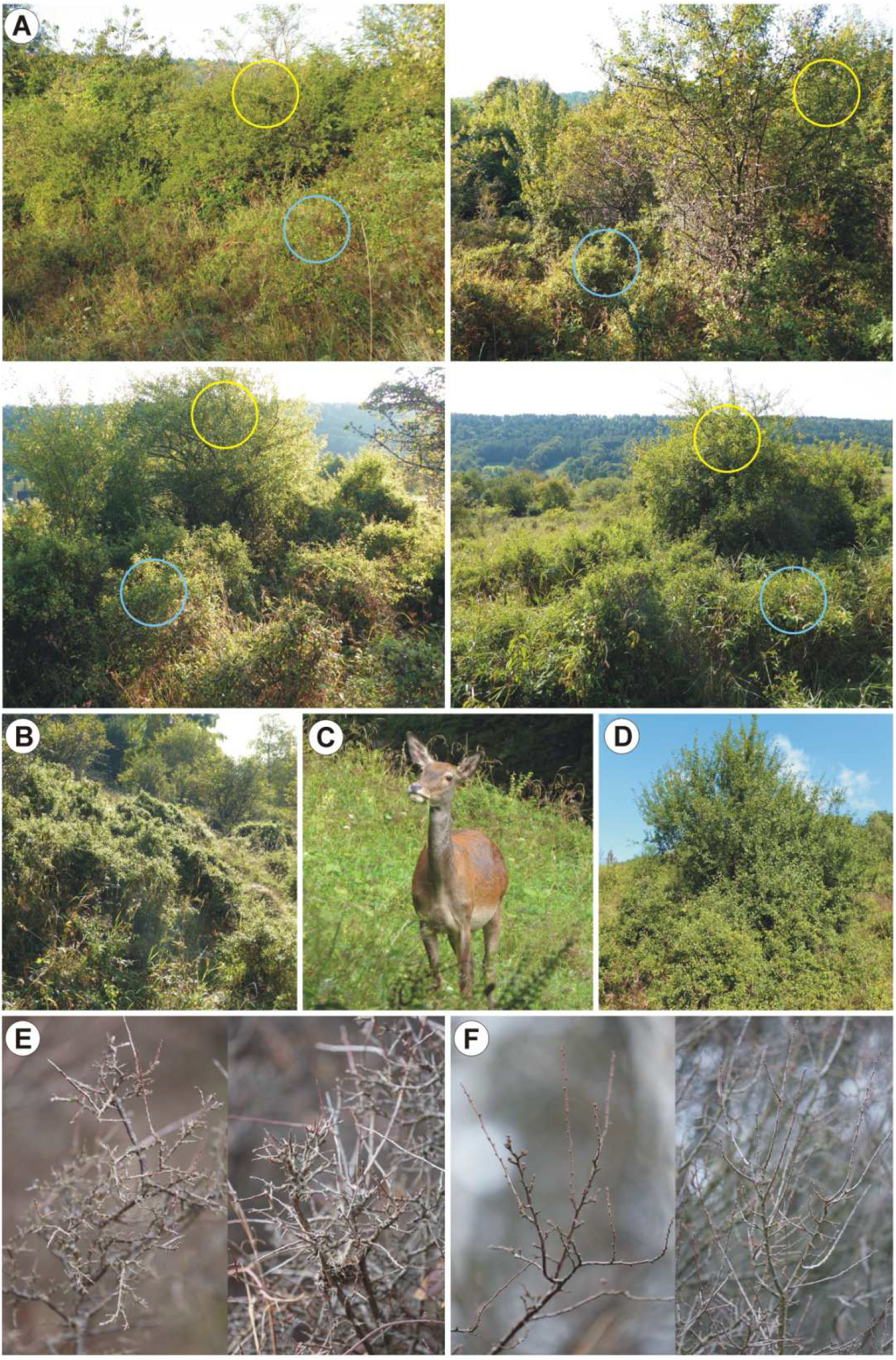
Blackthorn *Prunus spinosa* thickets subjected to long-term browsing by red deer *Cervus elaphus* (Bieszczady Mountains, SE Poland); A – examples of sites where paired samples were collected, yellow and blue circles indicate unbrowsed and browsed blackthorns, respectively; B – examples of heavily browsed blackthorns; C – female red deer; D – an unbrowsed blackthorn bush; E – typical top shoots of heavily browsed blackthorns; F – typical top shoots of unbrowsed blackthorns.

The study area has been routinely inspected over the past 20 years to evaluate the presence of ungulates; red deer regularly forage in it. During the growing season, from March to October, it was often visited by several deer, and herd sizes occasionally exceeded several dozen individuals (Fig. 1C). Single roe deer *Capreolus capreolus* were also sometimes recorded there. During the last 20 years, the area has not been grazed by livestock such as sheep, goats or cattle. Predators occurring there include the wolf *Canis lupus*, and domestic dogs occasionally roaming freely off the leash (people walk their dogs in the study area).

### 2.2. Field methods

To investigate for differences in chemical composition, 30 blackthorn bushes were randomly selected from those growing in the study area. In the vicinity of each unbrowsed bush, a browsed one was selected at random within a maximum radius of 5 m (Fig. 1A). During the selection, blackthorns to which access by herbivores was obstructed, i.e. unbrowsed bushes in the centres of extensive and dense clumps of shrubby vegetation or situated on the forest edge, were not included in the sampling. Approximately 250 g of top shoots, representing the current year’s growth, were taken from each individual in September 2000. Each pair of samples (one sample from a browsed bush and one from the paired unbrowsed bush) was deposited in a refrigerator (temperature ca 5°C) within a maximum of 2 h after collection, followed by a maximum of another 12 h in a freezer (–18°C) and another 12 h in a deep freeze (–80°C).

### 2.3. Chemical composition

Leaves were separated from twigs for the laboratory analyses. The leaf samples were dried and then ground in a knife mill (Retsch, GM 200). The C and N contents of the leaves were determined using an elemental analyser LECO CNS (TrueMac Analyzer; Leco, St. Joseph, MI, USA). The concentrations of macro- and microelements in the leaf samples were determined by inductively coupled plasma-optical emission spectrometry (ICP-OES Thermo iCAP 6500 DUO, Thermo Fisher Scientific, Cambridge, UK) after prior mineralization. Dried samples of leaves were mineralized in a mixture of HNO_3_ and HClO_4_ (3:1). The weight moisture was calculated using the following formula:

Mw = ((Mn-Md)/Mn) × 100 [%],

where: Mw – weight moisture, Mn – weight of fresh sample, Md – weight of dried sample.

### 2.4. Biochemical composition

Leaves detached from the shoots were rapidly frozen in liquid nitrogen and pulverized into a uniform powder, which was then weighed to determine the fresh weight. Subsequently, the samples were lyophilized (Labconco FreeZone 2.5 Liter Freeze Dry System, Kansas City, USA) for 120 h. The samples were then re-weighed to determine the dry weight and used for biochemical analysis.

#### 2.4.1. Photosynthetic pigments

Photosynthetic pigments were extracted according to Lichtenthaler and Welburn (1983) with modifications for a microplate reader. Approximately 15 mg of lyophilized leaf powder were extracted twice with methanol, the two extracts then being pooled. The absorbance of each sample was measured using a Synergy-2 microplate reader (Biotek, Winooski, VT, USA) at wavelengths of 666 nm, 653 nm and 470 nm. The concentrations of chlorophyll a (C_Chla_), chlorophyll b (C_Chlb_) and carotenoids (including xanthophylls and carotenes, C_x+c_) were calculated using the following formulas:

C_Chla_ = 15.65 × A_666_ – 7.34 × A_653_ [µg/ml],

C_Chlb_ = 27.05 × A_653_ – 11.21 × A_666_ [µg/ml],

C_x+c_ = (1000 × A_470_ – 2.86 × C_Chla_ – 129.2 × C_Chlb_)/245 [µg/ml].

#### 2.4.2. Soluble and insoluble proteins

Proteins were determined using the Bradford method (Bradford 1976) as modified by Ernst and Zorr (2010) for use with a microplate reader. Approximately 15 mg of lyophilized leaf powder were extracted for 1 h on ice with a buffer consisting of 100 mM potassium phosphate (pH 7.8), 2.0 mM ethylenediaminetetraacetic acid with 0.5% (w/v) Triton X and 1 mM dithiothreitol. The extract obtained after centrifugation was used to determine the content of soluble proteins, while the pellets were reused to extract and determine the content of insoluble proteins using a buffer containing SDS (0.05% w/v sodium dodecyl sulphate). To determine the soluble and insoluble protein contents, the previously obtained extracts were mixed with Bradford reagent and incubated in the dark for 5 minutes. The absorbance was measured at 450 and 590 nm using a Synergy-2 microplate reader (Biotek, Winooski, VT, USA). The protein content was calculated from the calibration curve using bovine serum albumin as standard.

#### 2.4.3. Soluble and reserve carbohydrates

The content of soluble carbohydrates was determined using the method of Dubois et al. (1951), adapted for a microplate reader by Marcińska et al. (2013), with minor modifications. Approximately 15 mg of lyophilized leaf powder were extracted with 80% ethanol for 30 minutes at 50°C. After centrifugation, the pellet was retained to determine starch, and the extract was used to determine soluble sugars. For this purpose, 200 µl of the diluted extract were mixed with 600 µl of conc. H_2_SO_4_ and 120 µl of 5% aqueous phenol solution, then incubated for 15 minutes at 90°C. The absorbance was read at 490 nm, and glucose was used as standard. The reserve carbohydrates were determined using the enzymatic method (Gomez et al. 2007). The pellet left after the extraction of soluble carbohydrates was resuspended in water and heated for 1 h at 98°C to dissolve the starch. Then, to each sample, amyloglucosidase (100 units) and α-amylase (70 units) in citrate buffer (pH 4.5) were added and the samples incubated at 56°C for 90 minutes. The resulting glucose was measured using hexokinase with glucose 6-phosphate dehydrogenase in the presence of ATP and NADP+ and determined spectrophotometrically at 340 nm. All the enzymes used for the carbohydrate analysis were manufactured by Megazyme International Ireland (Wicklow, Ireland). The contents of photosynthetic pigments, soluble and insoluble proteins, as well as soluble and reserve carbohydrates were expressed in mg g^-1^ (d.m.).

### 2.5. Data handling and analyses

The differences between the biochemical leaf profiles of the two groups of blackthorn (browsed *vs* unbrowsed) were inspected using the Mann-Whitney U test. Specifically, we checked the differences in the content of each variable: N (%), C (%), Ca (mg kg^-1^), cadmium Cd (mg kg^-1^), cobalt Co (mg kg^-1^), chromium Cr (mg kg^-1^), Cu (mg kg^-1^), iron Fe (mg kg^-1^), potassium K (mg kg^-1^), Mg (mg kg^-1^), manganese Mn (mg kg^-1^), sodium Na (mg kg^-1^), nickel Ni (mg kg^-1^), P (mg kg^-1^), lead Pb (mg kg^-1^), zinc Zn (mg kg^-1^), soluble carbohydrates (mg g^-1^ d.m.), reserve carbohydrates (mg g^-1^ d.m.), soluble proteins (mg g^-1^ d.m.), insoluble proteins (mg g^-1^ d.m.), chlorophyll a (mg g^-1^ d.m.), chlorophyll b (mg g^-1^ d.m.), carotenoids (mg g^-1^ d.m.) and water (%). To assess the relations between all the biochemical variables, we used Spearman’s correlation for all the samples pooled and separately for the browsed and unbrowsed groups. C:N, C:P, N:P and Ca:P ratios were calculated at the molecular level. The differences between the elemental ratios of the two groups of blackthorn (browsed *vs* unbrowsed) were inspected using the Mann-Whitney U test. Differences were assumed to be significant at a significance level of p < 0.05.

To investigate the differences between the two groups of blackthorns and their relationship with quantitative variables, we performed factor analysis of mixed data (FAMD). FAMD is a principal component method that combines principal component analysis (PCA) for continuous variables and multiple correspondence analysis (MCA) for categorical variables (Kassambara 2017). In our approach, all the variables describing the biochemical profile were treated as continuous, while Group, i.e. browsed *vs* unbrowsed blackthorns, was used as a categorical variable. The percentage of variances was explained by each principal component, and eigenvalues were visualized. We used R software, version 4.3.1 (R Core Team 2023) with the packages factoextra (Kassambara and Mundt 2020), ggplot2 (Wickham 2016), corrplot (Taiyun and Simko 2021) and ggpubr (Kassambara 2023).

## 3. Results

The two groups of blackthorns, i.e. browsed and unbrowsed, differed in the composition of 11 out of the 24 variables: C, N, P, Ca, Mg, Cu, carotenoids, water, chlorophyll a, chlorophyll b and insoluble proteins (Fig. 2). The browsed blackthorns had higher concentrations of 9 out of 11 nutrients, except for Ca and Mg, the values of which were lower than for the unbrowsed group (Fig. 2). The C:N, C:P, N:P and Ca:P ratios were higher in the unbrowsed than in the browsed group (Table 1). The correlation between C and P contents was positive in the browsed group (Fig. 3B) but negative in the unbrowsed one (Fig. 3C). Group (categorical variable) contributed to both principal dimensions (Dimension 1 = 4.6%; Dimension 2 = 11.1%) of FAMD (Fig. 4B-E). N, chlorophyll a, carotenoids, water, chlorophyll b, insoluble proteins, soluble proteins, reserve carbohydrates and K were represented in Dimension 1 (Fig. 4D), while Mg, P, Mn and Ca were represented in Dimension 2 (Fig. 4E).

**Fig. 2.**
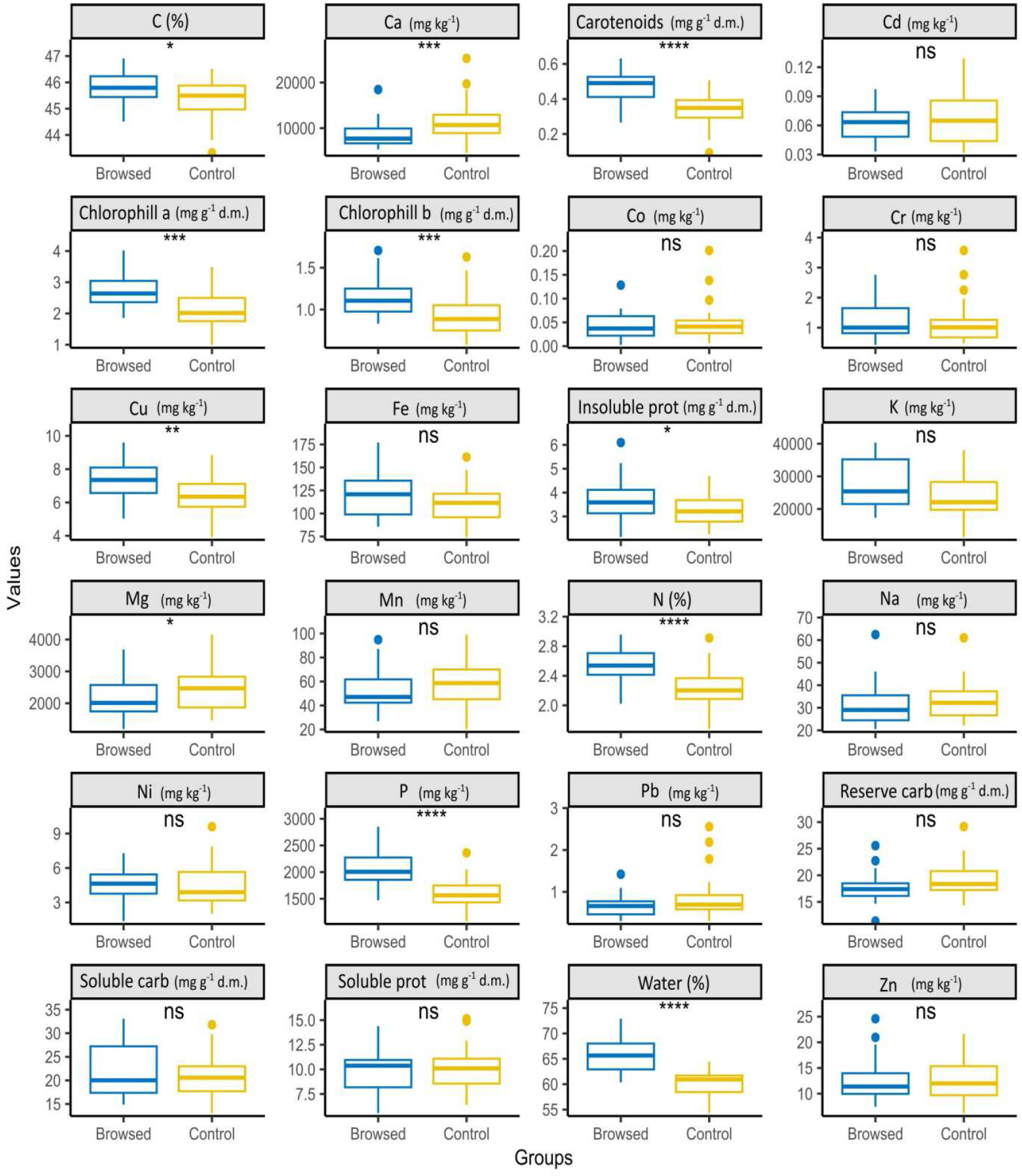
Differences between the concentrations of elements and nutrients present in browsed and unbrowsed (control) blackthorn *Prunus spinosa* leaves subjected to long-term pressure by red deer *Cervus elaphus*. The horizontal line, box, vertical line and points represent the median, quartile range, range and outliers, respectively. Differences were tested with the Mann-Whitney U test. Abbreviation: carb − carbohydrates, prot – proteins. The symbols refer to exact *p*-values: ns – p≥0.05; * – p<0.05; ** – p<0.01; *** – p<0.001; **** – p<0.0001, respectively.

**Fig. 3.**
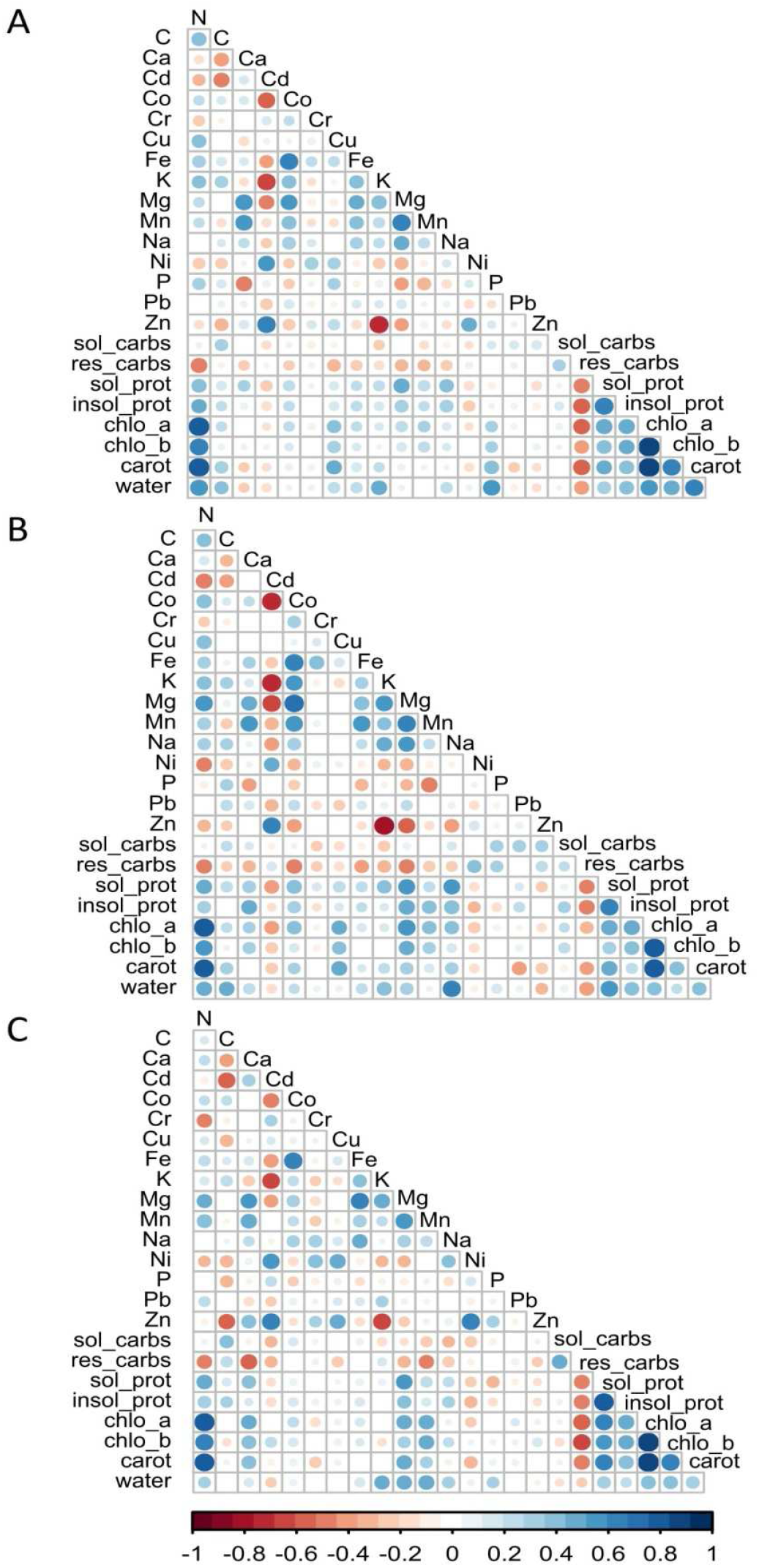
Spearman rank correlation matrix between all the variables identified in browsed and unbrowsed (control) leaves of blackthorns *Prunus spinosa* subjected to long-term pressure by red deer *Cervus elaphus*. A shows the relationships for all the data, while B and C present the relationships for browsed and unbrowsed blackthorns, respectively. The value of the correlation is proportional to the size of the inner circle and colour intensity (blue – positive, red – negative). Abbreviation: sol_carb – soluble carbohydrates, res_carb – reserve carbohydrates, sol_prot – soluble proteins, insol_prot – insoluble proteins, chlo_a – chlorophyll a, chlo_b – chlorophyll b, carot – carotenoids.

**Fig. 4.**
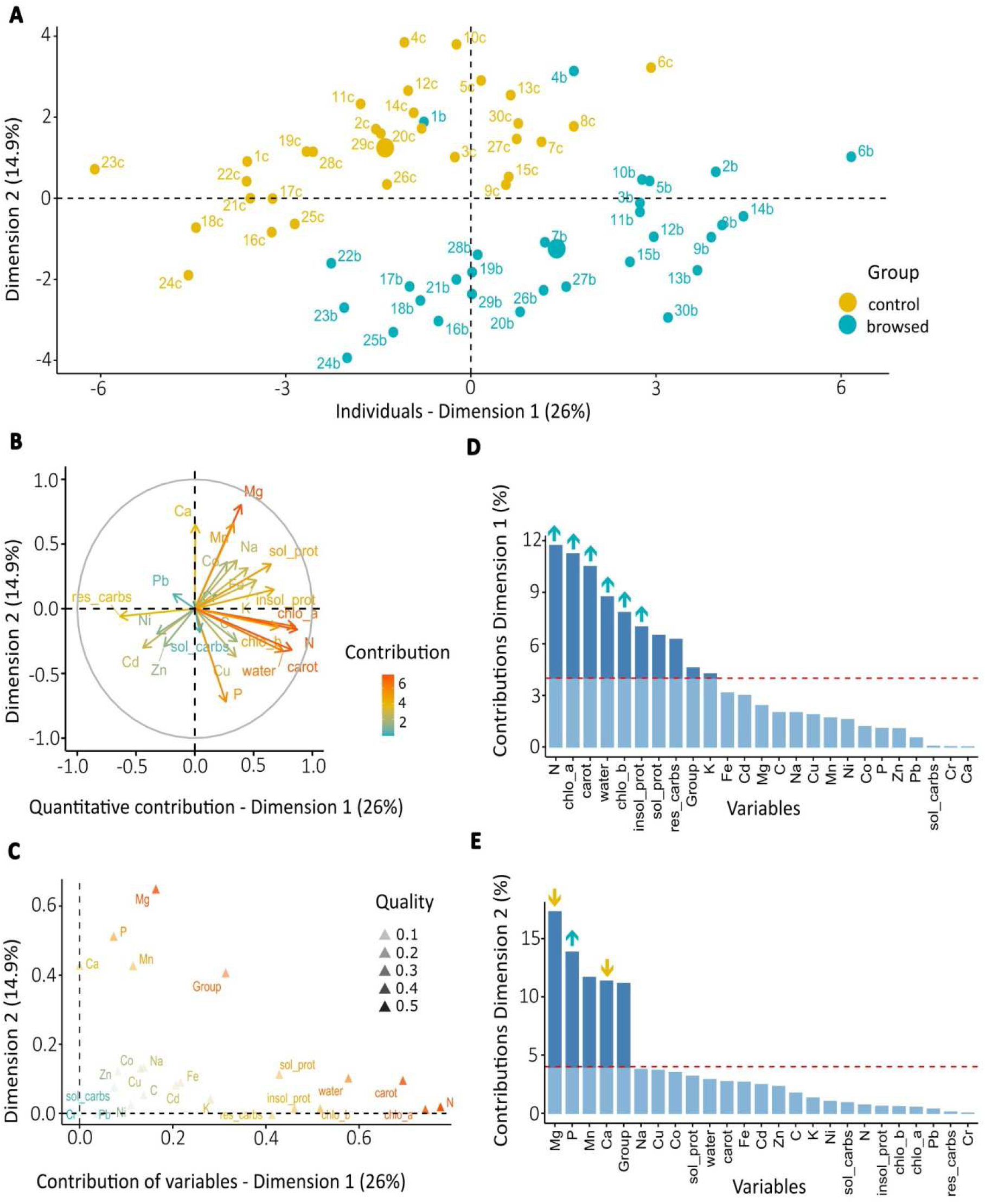
Differences in nutritional characteristics of blackthorn *Prunus spinosa* subjected to long-term pressure by red deer *Cervus elaphus* as revealed by factor analysis for mixed data (FAMD): A – plot for an individual blackthorn bush; each dot represents a single individual and each browsed bush denoted by the letter b is coupled with a control (unbrowsed) bush denoted by the letter c and associated with the same number; B – correlations of quantitative variables with both dimensions; the value of the contribution is defined by the gradual colouration (red = high, blue = low); variables pointing in the same direction are positively correlated; the length of the arrow indicates the description power of the variable in the dimensions; C – relationships between variables contributing to both dimensions; the value of the contribution is defined by gradual colouration as in B and the quality of representation for each variable is indicated by the transparency; D-E – contribution (%) of the variables to dimensions 1 (D) and 2 (E); each bar represents a variable; the red line shows the expected values if all the variables make equal contributions to the dataset; the area below the red line is shaded and the arrows indicate the difference between the browsed and unbrowsed groups (see Fig. 2) for each variable (up – median value higher in the browsed group, down – median value lower in the browsed group). For the abbreviations of the variables, see Fig. 3.

**Table 1.** Differences in the elemental ratios: Carbon:Nitrogen (C:N), Carbon:Phosphorus (C:P), Nitrogen:Phosphorus (N:P) and Calcium:Phosphorus (Ca:P) in leaves of blackthorn *Prunus spinosa* browsed and unbrowsed by red deer *Cervus elaphus*.

| Ratio | Browsed<br>median ratio | Unbrowsed<br>median ratio | W | *p* |
| --- | --- | --- | --- | --- |
| C:N | 21.1 | 24.1 | 177 | 0.000 |
| C:P | 585.3 | 737.7 | 116 | 0.000 |
| N:P | 27.4 | 31.4 | 282 | 0.013 |
| Ca:P | 2.8 | 5.3 | 139 | 0.000 |

### 3.1. Macroelements

#### 3.1.1. C and N

The browsed and unbrowsed blackthorns differed in their C and N contents (Fig. 2), and both these elements were positively correlated (Fig. 3). N was positively correlated with carotenoids and chlorophyll a, but negatively with reserve carbohydrates (Fig. 3). The correlation between C and water content was positive in the browsed group (Fig. 3B), but no such correlation was recorded in the unbrowsed one (Fig. 3C). N was also the variable making the largest contribution (11.7%) to the first dimension (Fig. 4D), whereas C was not represented in either dimension of FAMD (Fig. 4).

#### 3.1.2. Ca, Mg and P

The browsed group had lower concentrations of Ca and Mg and a higher concentration of P compared with the unbrowsed group (Fig. 2). Ca was negatively correlated with P but positively with both Mg and Mn. P was positively correlated with water and carotenoids, but negatively correlated with Mg, Mn and Na. Mg was positively correlated with Mn but negatively with Cu (Fig. 3). Mg (17.3%), P (13.8%) and Ca (11.4%) made a substantial contribution to the second dimension of FAMD (Fig. 4E).

#### 3.1.3. K and Na

K and Na did not differ between the two groups, although the K concentration approached (*p* = 0.055) the significance level, taking a marginally higher median value in the browsed group (Fig. 2). The correlation between K and Na contents was positive in the browsed group (Fig. 3B), but no such correlation was recorded in the unbrowsed group (Fig. 3C). K contributed to the first dimension of FAMD (4.2%), whereas Na made no contribution to either dimension (Fig. 4).

### 3.2. Microelements (Cu, Mn, Fe and Zn)

The browsed group had a higher concentration of Cu than the unbrowsed group (Fig. 2), whereas Mn, Fe and Zn did not differ between these groups. The Mn concentration was positively correlated with that of Mg (Fig. 3). The correlation between Mn and P contents was negative in the browsed group (Fig. 3B), but no such correlation was recorded in the unbrowsed group (Fig. 3C). Mn (11.7%) contributed to the second dimension of FAMD (Fig. 4E), but Cu, Fe and Zn did not contribute to either dimension (Fig. 4).

### 3.3. Water and biochemical elements

The contents of water and the biochemical elements were higher in the browsed group than in the unbrowsed group except for soluble carbohydrates, reserve carbohydrates (approaching significance level, *p* = 0.053) and soluble proteins (Fig. 2). Chlorophyll a (11.2%), carotenoids (10.5%), water (8.8%), chlorophyll b (7.8%), insoluble proteins (7.0%), soluble proteins (6.5%) and reserve carbohydrates (6.3%) contributed to the first dimension of FAMD (Fig. 4C). Proteins, chlorophylls, carotenoids and water were all positively correlated with each other (Fig. 3). Reserve carbohydrates were negatively or not correlated to all the other elements except for a weakly positive relationship with soluble carbohydrates (Fig. 3).

The contents of chlorophyll a, chlorophyll b and carotenoids correlated positively with the Cu concentration in the browsed group (Fig. 3B), but such correlations were not recorded in the unbrowsed group (Fig. 3C). Soluble carbohydrates and reserve carbohydrates contents were correlated positively with the P content in the browsed group (Fig. 3B), but such correlations were not recorded in the unbrowsed group (Fig. 3C). Reserve carbohydrates content correlated negatively with Co, Fe and K contents in the browsed group (Fig. 3B), whereas no such correlations were recorded in the unbrowsed group (Fig. 3C).

## 4. Discussion

The results of our analysis indicated distinct chemical profiles of blackthorn bushes subjected to long-term browsing by red deer in comparison with unbrowsed bushes. As suggested in other studies, the nutritional composition of foliage (structural carbohydrates, proteins, minerals) and the concentration of plant secondary compounds are drivers of ungulate browsing preferences (Ceacero et al. 2015; Holeski et al. 2016). Ceacero et al. (2015) state that red deer prefer feeding on shrubs with lower contents of Cu and Zn, below-deficiency levels of K, Mg, Na and P, while avoiding plants with surplus concentrations of S (max tolerance = 0.2%), Na (max tolerance = 0.35%) and Ca (max tolerance = 2%) (Suttle 2010; Ceacero et al. 2015).

### 4.1. Macroelements

As growth and functioning strongly depend on N (Epstein and Bloom 2005), plants have evolved to cope with its heterogenous availability in the soil, i.e. in inorganic and organic forms (Kraiser et al. 2011). Moreover, plants are able to display an ample developmental plasticity, for example, by modulating the root system architecture to ensure adequate N supplies (Robinson 1994). Morphological plant traits, such as the specific leaf area, are good predictors of plant growth rates and are positively correlated with leaf N content (Wright et al. 2010), although other traits, such as leaf C or P contents, are linked to plant performance (Westoby et al. 2002).

Our results showed that the C and N concentrations were higher in heavily browsed blackthorns, and previous studies on forage preferences yielded similar results (Gowda 1997; Danell et al. 2003; Fornara and Toit 2007; Scogings et al. 2012; Santos Pimenta et al. 2014; Wigley et al. 2015; Peschiutta et al. 2018; Nosko et al. 2020). The higher N and C contents make a plant more attractive to herbivores because of its enhanced nutritional value (Nosko et al. 2020). N is a macro-element essential for the synthesis of amino acids, which are required for the proper growth of tissues and enzymatic activity in animal organisms, including deer. With N-rich food, animals can synthesize amino acids more intensively during reactions taking place in the rumen (Bach et al. 2005). Scogings et al. (2011) showed that lower levels of defensive compounds are accompanied by high N levels. In deer exclusion experiments, Barrere et al. (2019) found such a trade-off between investment in growth by increasing the nutrient uptake and investment in defence, either by increasing chemical or structural resistance, and also that the leaf C:N ratio was a good predictor of plant defence against herbivory. Depending on the plant species, higher C:N ratios were solely associated with structural defence, i.e. by investment in cell-wall compounds such as lignin (Agrawal & Fishbein 2006; Barrere et al. 2019) or chemical defence, i.e. by investment in carbon-based compounds such as phenolics (Royer et al. 2013; Barrere et al. 2019), or both (Mondolot et al. 2008). Although we did not measure the quantity of secondary N-based defensive metabolites, e.g. cyanogenic glycosides such as amygdalin and prunasin (see section 4.4 – Study limitations), we found that the C:N ratio was higher in unbrowsed bushes. On the other hand, we observed that thorns were better developed in browsed than in unbrowsed blackthorns, which could indicate that the higher C:N ratio in the unbrowsed group reflected chemical defence only.

Savanna species can be categorized into groups in accordance with their defence strategies (Wigley et al. 2018). One such group contains species exhibiting a high nutritional status and a well-developed form of structural defence, such as spines. Another group contains species of low nutritional quality and a high level of chemical defence. In our study, a high C:N ratio indicated possible chemical defence in unbrowsed individuals in conjunction with low nutritional status. Thus, we may expect that a mechanism similar to the one recorded in savanna species may be playing a part in between-individual variation of the same species subjected to herbivory. We recorded the better development of thorns in browsed blackthorns, which could have resulted from shoot modifications as a response to constant yearly browsing. Thus, the mechanism of blackthorn defence to herbivory that we observed may be complex and will need further investigation.

In our study, both Ca and P were important contributors to the bushes in the unbrowsed group. P is a component of key molecules such as nucleic acids, phospholipids and ATP (Vitousek et al. 2010). The form of P most readily accessed by plants is inorganic phosphate (Pi), which is absorbed by the roots and transported to the young leaves (Mimura et al. 1996).

However, there is also significant retranslocation of Pi from old leaves to the growing shoots and from the shoots to the roots. Moreover, N:P ratios are negatively correlated with the relative growth of a species and its N-indicator values (Güsewell 2004). Ca is an essential plant nutrient playing a structural role in membranes and cell walls, also acting as a messenger in physiological processes and a regulator responding to environmental stresses and growth-related processes (White and Broadley 2003, Lautner & Fromm 2010). Environmental and physical stress in plants is known to produce Ca impulses and to trigger Ca-dependent electrical signals within the plant (Fromm & Spanswick 1993; Lautner & Fromm 2010). Even if Ca deficiency seems rare in nature, the Ca shoot concentration in optimal conditions ranges from 0.1 to 5% of dry mass (White and Broadley 2003; Ceacero et al. 2015). Consequently, the higher value of Ca found in unbrowsed blackthorns suggests a potential role of Ca-based defences influencing browsing selectivity (Ward et al. 1997; Molano-Flores 2001; Ruiz et al. 2002; Hanley et al. 2007; Ceacero et al. 2015; Peschiutta et al. 2020). Calcium oxalate crystals (COC) are efficient against invertebrate herbivory, but their presence in all plants suggest that they also use it to deter mammalian herbivores (Ward et al. 1997; Molano-Flores 2001; Ruiz et al. 2002; Peschiutta et al. 2020).

As expected, because of the physiological mechanisms that see Ca and P as antagonists (Benge 2002; Miller et al. 2003), our results confirmed a strong negative correlation between these elements. We found the Ca:P ratio to be lower in the browsed than in the unbrowsed group (see Table 1). The optimal Ca:P ratio for ungulates was found to be close to 1:1 or 2:1 (Ceacero et al. 2015; Grzegorczyk et al. 2017), even if ruminants can tolerate a ratio of up to 6:1 (Ca max tolerance = 2%; P max tolerance = 0.6%) (Suttle 2010; Ceacero et al. 2015). Above this maximum ratio, their interaction will reduce the absorption of P, leading to a negative effect on the ungulate metabolism (Benge 2002; Miller et al. 2003). Compared with the results of Wigley et al. (2015) on acacia *Acacia* sp., our browsed blackthorns had a higher P concentration. P is considered, together with N, to be the main driver of plant stoichiometry (Ågren and Weih 2020). However, with our methodology we were unable to determine which effect was the driving force: was the value of P in browsed individuals higher because of the browsing itself or was it a consequence of a physiological interaction that optimizes the choice for ungulates?

The lower concentration of Mg is responsible for the structural changes of the ribosomes and reduces rates of photosynthesis (Mehne-Jakobs, 1996). This element is also known to be essential for muscle contraction and energy metabolism in mammals (Miller et al. 2003; Ceacero et al. 2015; Hollingsworth et al. 2021). Our results indicated that the Mg content was lower in the browsed blackthorns and that Mg was the element making the greatest contribution to the dimension representing the unbrowsed bushes. These results partially contrast with those in the literature, where it is reported that deer prefer plants with a below-deficiency level of Mg (Suttle 2010; Ceacero et al. 2015). Analysis of 191 feed plant species for sika deer *Cervus nippon* revealed that all plant species can provide sufficient amounts of Mg, which is not the case for Na and Ca (Mori et al. 2023). Although plants are an important source of Mg as regards meeting herbivore needs, the mechanism behind Mg bioavailability still needs to be identified (Wilkinson et al. 1990). Whereas other studies have shown that deer select plants with lower Na and K contents (Suttle 2010; Ceacero et al. 2015), the difference in the contents of these elements was not significant in our analyses. Na is not represented by the principal dimensions of FAMD, and the contribution of K was minimal.

### 4.2. Microelements

Far fewer authors have examined the role of micronutrients in the development and growth of trees, essential though they are, than macronutrients (Hagen-Thorn and Stjernquist 2005). Fe and Mn are important for photosynthetic activity, and Cu participates in photosynthetic electron transport, mitochondrial respiration, cell wall metabolism and hormone signalling (Marschner 1995). On the other hand, Zn is an important component of proteins in plants, regulating their capacity for water uptake and transport (Kasim 2007) and reducing the adverse effects of short-term heat stress (Peck and McDonald, 2010). Our results indicate that the Cu content is higher in the browsed blackthorns. In mammals, Cu is essential for the proper growth of skeletal mass and in their metabolism (Miller et al. 2003; Hollingsworth et al. 2021). These results partially contradict the literature, where deer have been shown to prefer plants with lower Cu levels (Suttle 2010; Ceacero et al. 2015). However, as Cu is not represented in the FAMD, we assume that the effect of Cu is unlikely to make a significant contribution to the dietary preferences of deer.

The Fe concentrations in both the browsed and unbrowsed blackthorns lie within the optimal range suggested for ungulates, whereas the Zn level in both browsed and unbrowsed plants was below the required range for ungulates (Suttle 2010; Ceacero et al. 2015; Hollingsworth et al. 2021). We did not detect any difference between the two groups as regards the concentrations of these two elements. Moreover, Fe and Zn were not represented by the principal dimensions of FAMD, and their contribution to the variables’ representation was minimal (< 0.2%), which suggests that their role in deer foraging preferences is limited. However, Mn was the third element with the highest contribution to the second dimension of FAMD, indicating the potential importance of this element in the preferences of individual blackthorns.

### 4.3. Water and biochemical elements

Intense browsing affects the photosynthetic capacity of an entire plant (Peschiutta et al. 2018), and high concentrations of insoluble proteins, chlorophylls, carotenoids and water in browsed individuals support the hypothesis that there is a response to herbivory-induced stress (Peschiutta et al. 2020). Higher levels of assimilation pigments and carotenoids in a browsed plant may indicate its attempt to compensate for losses incurred due to foraging. A similar response was observed, for example, during aphid feeding on the assimilation apparatus of wheat, leading to increased levels of chlorophyll a, b and carotenoids in uninfested tissues adjacent to those damaged by aphids (Ni et al. 2002). Enhanced photosynthetic activities may increase the growth rate and induce changes in the allocation and distribution of nutrients within the plant (Gordon and Prins 2008; Orians et al. 2011; Scogings et al. 2014; Peschiutta et al. 2018). The greater content of insoluble proteins (mainly membrane and trans-membrane proteins) in browsed than in unbrowsed plants signifies the greater activity of the enzymatic apparatus involved in the production of defensive substances (protein and non-protein) and enhances trans-membrane transport and intercellular signalling. Despite carbohydrates being the leading nutritional components of the ungulate diet (Holeski et al. 2016), we found no difference in their levels between browsed and unbrowsed trees. Moreover, reserve carbohydrates play only a limited role in the characterization of the browsed blackthorns. On the other hand, previous studies (Ceacero et al. 2015; Holeski et al. 2016) have shown that proteins may mediate feeding preferences.

### 4.4. Study limitations

The results concerning the main relevance of defence mechanisms (chemical – morphological) against long-term browsing are inconsistent (Rooke et al. 2004; Wigley et al. 2015). Even if secondary metabolites are regarded as essential in herbivore-plant interactions (Foley and Moore 2005; Iason and Villalba 2006; Gordon and Prins 2008; Scogings et al. 2011; Perkovich and Ward 2022), it has been suggested that severe browsing can reduce the concentrations of some secondary metabolites, like tannins, in exchange for an investment in C-based defences (Pardo et al. 2018). Moreover, some individual plants may invest in higher growth rates rather than defence components based on the levels of available resources (Scogings et al. 2011; Pardo et al. 2018). Also, the structural response of a particular plant to ungulate browsing may vary to such an extent that some parts of the tree will be readily eaten, while other parts will be less attractive (Hartley et al. 1997); furthermore, the differences within one tree may be greater than those between two separate trees. On the other hand, the blackthorn’s spinescences are part of a well-known structural defence mechanism used against biotic stress factors (Pisani and Distel 1998; Scogings et al. 2011; Salek et al. 2019). We noticed that all heavily browsed blackthorns had thick, hard and well-developed thorns located on twigs of the external parts of the crowns, whereas on unbrowsed bushes such structures were less developed only on older twigs situated in the inner regions of the plants.

With our methodology we were unable to investigate whether herbivory was the reason for the chemical variability, i.e. whether a plant’s reaction was induced by browsing, or whether the inherited chemical variability of an individual blackthorn tree was indeed the reason driving food selection, i.e. initiating and sustaining the preferences/avoidances of foraging ungulates. There is evidence to show that the chemical response of vegetation can influence the foraging preferences of herbivores and mould between-tree selection (Gill 2001; Provenza et al. 2003; Scogings et al. 2012), and it has been suggested that the formation of a positive feedback towards already-browsed individuals may be a predictor of which individuals will be browsed again (Stolter 2008; Jager et al. 2009; Nosko et al. 2020). The high nutritional state can start a positive feedback loop where individuals already subjected to browsing are more likely to be browsed again (Stolter 2008; Jager et al. 2009). In the present case, we can speculate that the initial food selection was a random process, and that the influence of long-term browsing became apparent in the blackthorn’s morphology and chemical composition over time (see Vourc’h et al. 2002 for a similar study on browsing effects on western red cedar). Presumably, browsing pressure, initially random, mutates later into a more selective choice (Gordon and Prins 2008; Call and St. Clair 2018) driven by the difference in nutritional quality. However, there is still no clear explanation for this process and its effects on populations subjected to long-term browsing pressure.

## 5. Conclusions

Complex plant defence strategies combined with ungulates’ foraging preferences may contribute to plant-herbivore interactions. Our results revealed differences in the chemical and biochemical compositions of browsed and unbrowsed blackthorns forming a local population subject to long-term browsing by red deer. With two possible mechanisms mediating the observed chemical profile, i.e. the inherited initial profile or the effect of long-term browsing, we were able to identify clear differences in the nutritional characteristics of browsed and unbrowsed blackthorns. Our findings make a contribution to the knowledge of deer foraging preferences and plant adaptations to herbivore pressure in temperate ecosystems, and highlight the potential role of individual plant characteristics in the food selection process. The results are also helpful in understanding the evolution of plant-herbivore systems and vegetation dynamics, where a certain fraction of the population of a given plant species survives over the long term, despite severe pressure on the part of herbivores.

## Acknowledgements

This study was financially supported by the National Science Centre, Poland (grant No. 2019/03/X/NZ9/02005). Michał Ciach was supported by grant No. 2021/41/B/NZ8/03456 from the National Science Centre, Poland.

## Ethical statement

The study was performed in accordance with Polish law.

## Conflict of interest

The authors declare that they have no conflict of interest.

## Notes

### Competing Interest Statement

The authors have declared no competing interest.

